# FASTAFS: file system virtualisation of random access compressed FASTA files

**DOI:** 10.1101/2020.11.11.377689

**Authors:** Youri Hoogstrate, Guido Jenster, Harmen J. G. van de Werken

## Abstract

**Background:** The FASTA file format used to store polymeric sequence data has become a bioinformatics file standard used for decades. The relatively large files require additional files beyond the scope of the original format, to identify sequences and provide random access. Currently, multiple compressors have been developed to archive FASTA files back and forth, but these lack direct access to targeted content or metadata of the archive. Moreover, these solutions are not directly backwards compatible to FASTA files, resulting in limited software integration.

**Results:** We designed linux based a toolkit using Filesystem in Userspace (FUSE) that virtualises the content of DNA, RNA and protein FASTA archives into the filesystem. This guarantees in-sync virtualised metadata files and offers fast random-access decompression using Zstandard (zstd). The toolkit, FASTAFS, can track all system wide running instances, allows file integrity verification and can provide, instantly, scriptable access to sequence files and is easy to use and deploy.

**Conclusions:** FASTAFS is a user-friendly and easy to deploy backwards compatible generic purpose solution to store and access compressed FASTA files, since it offers file system access to FASTA files as well as in-sync metadata files through file virtualisation. Using virtual filesystems as in-between layer offers the possibility to design format conversion without the need to rewrite code into different languages while preserving compatibility.

**Code Availability:** https://github.com/yhoogstrate/fastafs

## Background

FASTA is a file format used for storing nucleotide and amino acid polymeric sequences and is compatible with a high variety of bioinformatics software. It is used as database for ribosomal RNA sequences, but also for eukaryotic reference genomes and protein databases that can be several gigabytes in size. In contrast to for example GenBank, it offers very limited support for metadata. Corresponding fai-index files are used to achieve random access by providing the sequence length, padding corrected file positions and padding and line length. This is static information that is embedded in the FASTA file, which is extracted after generating the FASTA file.

Scientific demand for reproducibility and interoperability of both software applications and data is growing strongly and as a result unique identification and data integrity play a critical role. In the CRAM data format, for instance, Next Generation Sequence (NGS) alignments are compressed relative to a reference sequence. In this format, the reference sequences are addressed using their unique identifier for interoperability. With the identifier, the corresponding sequence can be obtained directly using the online European Nucleotide Archive (ENA) service (https://www.ebi.ac.uk/ena/cram/swagger-ui.html), preserving the intrinsic link between the data file and the reference sequences. Because real-time computation of identifiers can be computationally expensive, they are stored in separate dictionary files (*.dict). Dict-files are, like fai-index files, beyond the scope of the original file format and have to be generated and maintained after obtaining the FASTA file.

Current software applications make use of FASTA files as input in two different manners:

- First, a tool reads a FASTA text file sequentially and in one-direction, starting with the first character in the file. For example, short-read alignment algorithms, but also motif-scanners that iteratively search for a given motif [1] across a sequence, read a FASTA file sequentially into the memory before building an index [2], [3]. Similarly, Single-Nucleotide Polymorphism (SNP) detectors may read by iterating sequentially over a FASTA file [4].
- Second, a tool reads a FASTA file in a random-access fashion by starting at an arbitrary location in the file and has the possibility to make jumps, forwards but also backwards, through the file. The precise file coordinates is typically calculated using the fai-index file. For example, a request to a genomic region within a genome browser is such a *random-access* request, since a next query can be expected at any genomic location. If underlying FASTA file access does not support jumping through a file it is necessary to copy a file entirely into memory. This procedure is extremely resource intensive and can slow a process significantly. Bioinformatics tools that rely on random-access in FASTA files are for i.e. JBrowse [5], samtools mpileup for VarScan2 [6]. But tools for quick file operations such as SeqKit [7], GATK [8] and Picard [9] also rely on random-access, of which the latter two require dict-files as well.

## Compression

The simplicity of the FASTA format makes the format convenient to work with. The trade-off is the requirement of the additional fai-index and dict-files, as well as having a relatively large file size. The large file size issue has been tackled by various compression methods [10], [11]. Although modern compressors achieve high compression ratios, most bioinformatics applications that require FASTA files are only rarely compatible with compressed equivalents. The only exception is occasional compatibility with gzipped FASTA.

Sequence compression algorithms create a compressed file (archive) yielding the compressed content. To use the original data, the archive needs to be fully decompressed into a temporary FASTA file again, unless the decompression algorithm also provides an Application Programming Interface (API) in the desired programming language. For instance, short read compressor DNA Sequence Reads Compressor 2 (DSRC2) [12] provides an API in C, C++ and Python.

The index algorithm of RNA read aligner The Spliced Transcripts Alignment to a Reference (STAR) [3] can be provided with the path to any decompression binary as argument and thus offers a generic solution to provide on-demand de-compression. However, implementing a similar solution in other applications would only work for applications with streaming instead of random access to FASTA Files. An analogues workaround to avoid file duplication is to make use of (named) pipes [10]. A pipe is a virtual, one-directional, data stream, that stays in idle as long as no further data requests come in. This could e.g. be the output of a decompressor. This is resource efficient as data access is chunked, but is not a generic solution as it does not offer random access. Access to FASTA archives in a random-access use case requires an available compression API that supports random access explicitly. If these conditions are not met, the primary goal of compression is then in practice lost. The FASTA file is still needed and having both the original and its compressed equivalent costs effectively more space rather than it saves.

Currently available bioinformatics applications that make use of FASTA files in a random-access setting mostly support only FASTA files and no compressed equivalents. Therefore, it is in practice necessary to keep a flat copy of a FASTA file with the corresponding the fai-index file. For systems limited to applications with streaming access to FASTA files, a decompression binary in combination with (named) pipes is an ideal way to use FASTA archives, although it requires management of metadata files. Instead of using a classical file converter binary for decompression, we can also file virtualisation. This way, file virtualization functions as layer between a compressed archive and the virtually mounted FASTA plus metadata files, which offers multiple advantages over classical (de-)compression binaries:

- Virtual files and their system calls are identical to flat file system calls. For tools that are only compatible with FASTA files, this preserves backwards compatibility, also for random access use-cases.
- There is no need to use additional disk space for temporary decompression and no need to read entire FASTA files into memory.
- For random access requests, computational resources are only spent on decompressing the region of interest.
- Implementations of compression and decompression in other programming languages or within other software applications are not needed, as it is backwards compatible with flat FASTA files.
- The archive is guaranteed to provide dict- and fai-index files that are in sync with their FASTA file of origin. This makes additional management of these metadata files unnecessary.

Making use of virtualization as layer between archive and decompressed content is a generic purpose solution as it provides random access to the original files. However, random access compression algorithms have typically smaller compression ratios. Moreover, maintaining virtual mount points requires effort at system administration level, for which FASTAFS provides a solution in its feature-rich toolkit. Here, we propose FASTAFS, a file archival format and toolkit that allows file integrity verification and provides unique sequence identifiers. In addition, it virtualises FASTA and guaranteed in-sync dict- and fai-index files files, from compressed 2-4 or 5-bit encodings.

## Implementation

FASTA File System (FASTAFS) file format consists of four blocks including (*1*) File Header (*2*) Per-Sequence-Data (*3*) Per-Sequence-Header and (*4*) File Metadata, to efficiently store sequence and metadata (**Figure 1**). During conversion, the metadata flag sets the archives status to incomplete. Each block of compressed sequence data is followed by the CRAM format and BAM specification compatible MD5 checksum [13], [14]. In the last phase of file conversion, file pointers are put in place and a metadata flag is updated to mark the archives conversion status to complete. The file ends with the CRC32 checksum used for whole file integrity verification.

**Figure 1:**
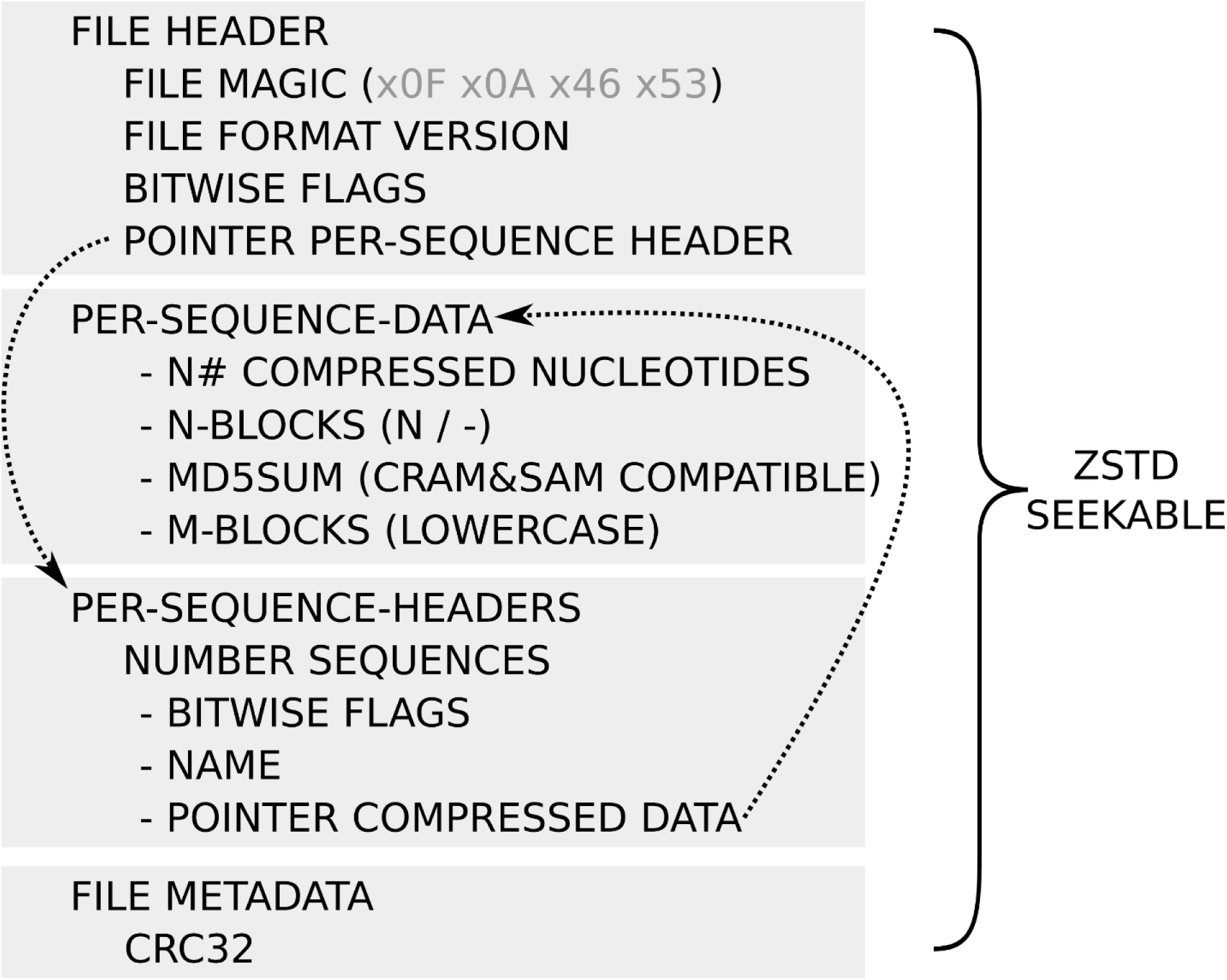
Overview of the FASTAFS file format specification. The layout of the FASTAFS format consists of four blocks, starting with the file header, followed by the per-sequence data, the per-sequence header data and a metadata block. The file header has a file pointer to the per-sequence header block, where each sequence has a file pointer to its data. The file ends with a metadata block, currently supporting a CRC32 checksum. The raw FASTAFS file is subsequently compressed with zstd-seekable. The full specification is available on the website: https://github.com/yhoogstrate/fastafs/blob/master/doc/FASTAFS-FORMAT-SPECIFICATION.md.

Sequence compressor Nucleotide Archival Format (NAF) [10] compresses sequence data first with a 4-bit encoding followed by generic compressor Zstandard (zstd), but it lacks random access. Given that NAF achieves high compression ratios [10], FASTAFS was designed in a somewhat similar fashion as it first compresses sequence data to a lower bit encoding (2-bit, 4-bit or 5-bit), followed by the random-access implementation of zstd called zstd-seekable.

### FASTAFS Toolkit

The LINUX based FATSTA FS toolkit is a single executable (`fastafs`) with different subcommands. The package comes also with an executable ‘`mount.fastafs`’ to mount via command line or directly using the `/etc/fstab` table.

#### Cache

FASTA files can be converted to a FASTFS archive the ‘fastafs cache’ subcommand, which adds a reference to the FASTAFS file into a config-file (**Figure S1A**).

#### Mount

The ‘fastafs mount’ subcommand is used to mount a FASTAFS archive to a directory (mount point) to virtualise the FASTA, fai-index, dict and UCSC TwoBit files (**Figure S1A**). All files are mounted read-only. Mount points can be configured in `/etc/fstab` which requires using the binary *mount.fastafs* instead of the binary *fastafs*. These entries can be configured to automatically mount during boot. Upon a file request, the kernel requests, through the Filesystem in Userspace (FUSE), the FASTAFS toolkit to provide either file attributes such a timestamps, size or permissions, or to copy real-time decompressed file content into a buffer.

In addition, FASTAFS provides filesystem access to query partial sequences using a subsequence identifier as filename in the ‘seq’ subdirectory. For example, the file `<mountpoint>/seq/chr1:10-20`contains only the sequence of this region, without additional characters such as newlines or spaces. Subsequently, requesting the file size of `<mount point>/seq/chr1` will provide its size in nucleotides. Indeed, these additional features do not solve backwards compatibility issues, but provide virtualised random access, without using the fai-index file, by functioning programming language independent API implemented at filesystem level.

#### List

The ‘fastafs list’ command gives an overview of the FASTAFS archives, their alias, number of sequences, format, compression ratio and all active mount points (**Figure S1A**).

#### View

Besides mounting, the FASTA contents can be decompressed to *stdout* using ‘fastafs view’, of which the padding can be set to a desired value and masking can be virtually disabled. The contents can also be exported to UCSC TwoBit format (**Figure S1B**).

#### Info

The ‘fastafs info’ subcommand gives information about the file layout, sequence size, the per-sequence MD5 checksum and used compression type. This subcommand can also be used to query European Nucleotide Archive (ENA) [15] whether the existence of a sequence MD5 checksum can be verified (**Figure S1C**).

#### Check

The ‘fastafs check’ command checks the file integrity using a CRC32 checksum. Integrity of compressed sequence data blocks can be checked separately using their MD5 checksums with the ‘--md5’ argument (**Figure S1D**).

#### ps

A list of active FASTAFS mount-points and their processes is provided by the ‘*fastafs ps’* subcommand. The mount point has an extended file attribute (xattr) named ‘FASTAFS-file’ that returns the mounted FASTAFS archive. When a FASTAFS file is mounted to multiple mount-points, they are each listed as separate entry with the corresponding system process id (**Figure S1E**).

FASTAFS format specification, toolkit and GPL-2.0 licensed C++ code is available at: https://github.com/yhoogstrate/fastafs

## Results

We compared the compression ratios of NAF, bgzip and MFCompress with FASTAFS (**Figure 2**). FASTAFS compression ratios for FASTA files with relatively few sequences (human reference genome: **GRCh38**, SARS-CoV-2 genome primary assembly (RNA): **NC_045512**.**2**, Coliphage phi-X174, complete genome **NC_001422**, fungus Neurospora crassa genome reference: **CM002240**) were similar as the ratios of NAF and MFCompress but not superior. For sequences with a relatively high number of sequences (miRNA, tRNA or protein databases), compression ratios of FASTAFS files are typically smaller than the other compressors, in particular for miRbase [16]. These files are composed of small sequences which result in a substantial contribution of the sequence names and MD5 checksums to the total archive file size.When the size of the archives is corrected with the space needed to store the MD5 checksums, the FASTAFS compression ratios are similar to those of MFCompress and NAF. Except for protein sequence compression, the most commonly used FASTA compression method (gzip) has consistently lower compression ratios than all other compressors.

**Figure 2:**
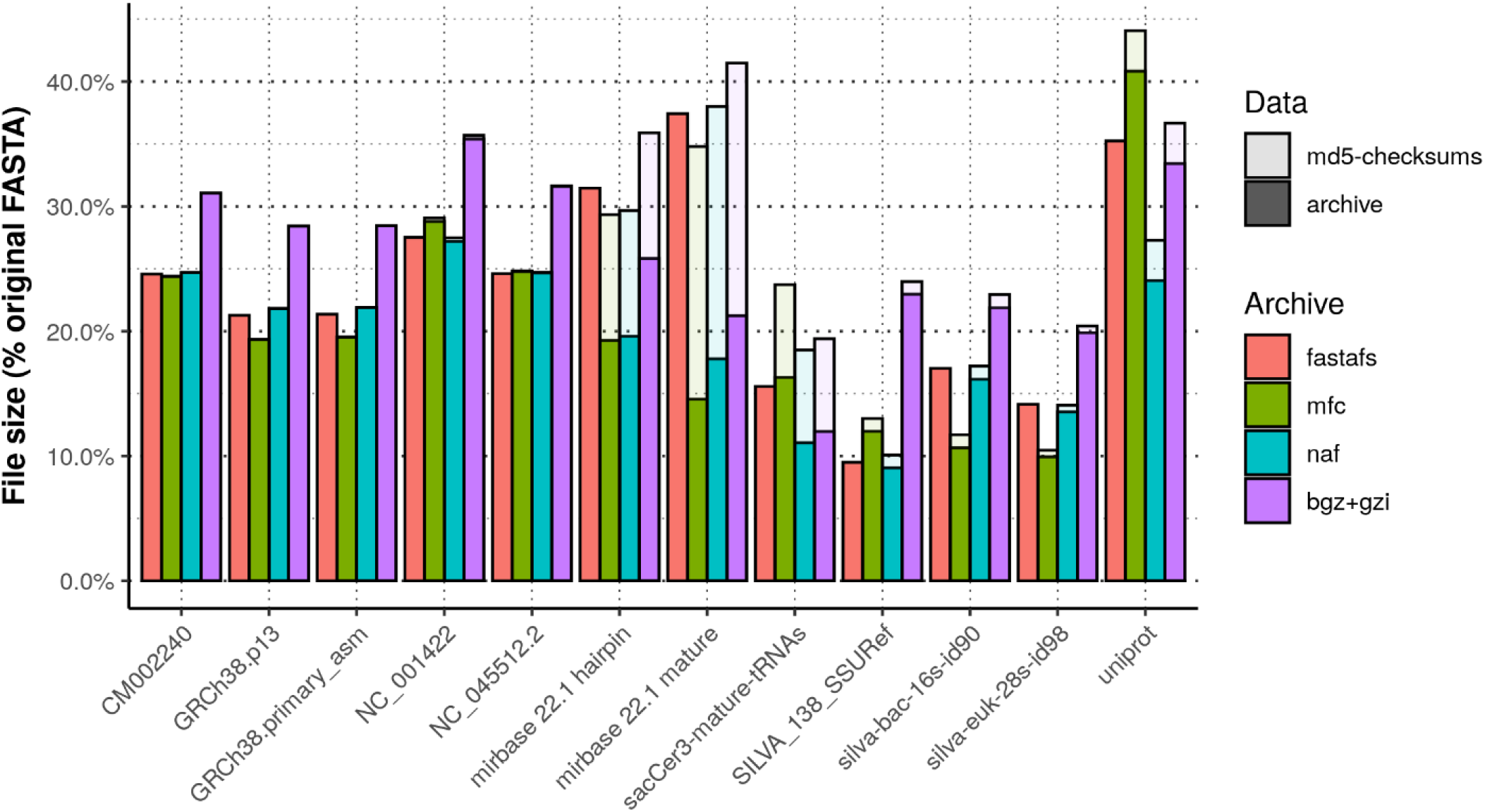
Overview of different archived files sizes. Comparison of compression ratios of a diverse set of FASTA files compressed with bgzip, MFCompress, NAF and FASTAFS. The bar height represents the percentage of the archives file size compared with the original FASTA files size. The translucent bars on top of the coloured bars represent the corrected file size needed to store 16 additional bytes per-sequence reserved for storing md5 checksums. We used genome references from fungi (CM002240), human with and without alternate loci (GRCh38.p13 and GRCh38.primary_asm), DNA (Coliphage phi-X174: NC_001422) and RNA viruses (SARS-CoV-2: NC_045512.2), databases with small RNAs (miRbase and tRNAs), Silva rRNA databases [20] and uniprot [21] for protein sequences.

## Conclusions

The FASTA file format is used to store biological polymeric sequence data in an easy-to-use format that has become a file standard in bioinformatics. Static information is embedded within each file, but needs to be extracted and stored in additional files to complement the FASTA file. We have developed a method, FASTAFS, to virtualise FASTA files along with their metadata files into the file system. The implementation makes use of the zstd-seekable compression library, which makes random access to the virtual FASTA files possible. FASTAFS comes with a feature rich toolkit that can manage the archives, their locations, their file integrity and provides file access in a backwards compatible manner to regular FASTA file access. This allows the archives to be used in existing software without the need for adaptation for compatibility and without the use of additional APIs.

Ideally, new bioinformatics analysis projects are started with a new folder that is under version control. This will allow the researcher to integrate FASTAFS with workflow management systems such as Snakemake [17] or Nextflow [18] as well as software dependencies by including dependency management configurations. Ultimately, this makes a project portable as it allows users to distribute projects over multiple locations, share it with other researchers and roll back to previous versions. Currently, version control for plain FASTA files is inconvenient and redundancy across multiple projects will occur quickly. However, by integrating FASTAFS mount points and scripts into a workflow management system FASTA files can be integrated intuitively into a projects’ version control. FASTAFS archives are currently compressed with a 2-bit, 4-bit or 5-bit encoding, followed by zstd-seekable, resulting in comparable compression ratios to other known compressors. Because the zstd-seekable implementation is still work in progress, adding additional free open source alternatives supporting random access such as bgzip [19] may be a future feature. FASTFS currently works with per-file aliases and CRAM compatible per-sequence identifiers. It would be more convenient to integrate FASTA files into workflow managers by using persistent per-file identifiers combined with a mechanism for decentralised synchronization of archives. As such additional features would be helpful; defining a system for per-file identifiers and development of decentralised file synchronization prompts future work. Overall, FASTFS is modern and elegant software solution for a user-friendly and easy to deploy generic purpose solution to store and access to compressed FASTA files.

---

**Figure S1A:**
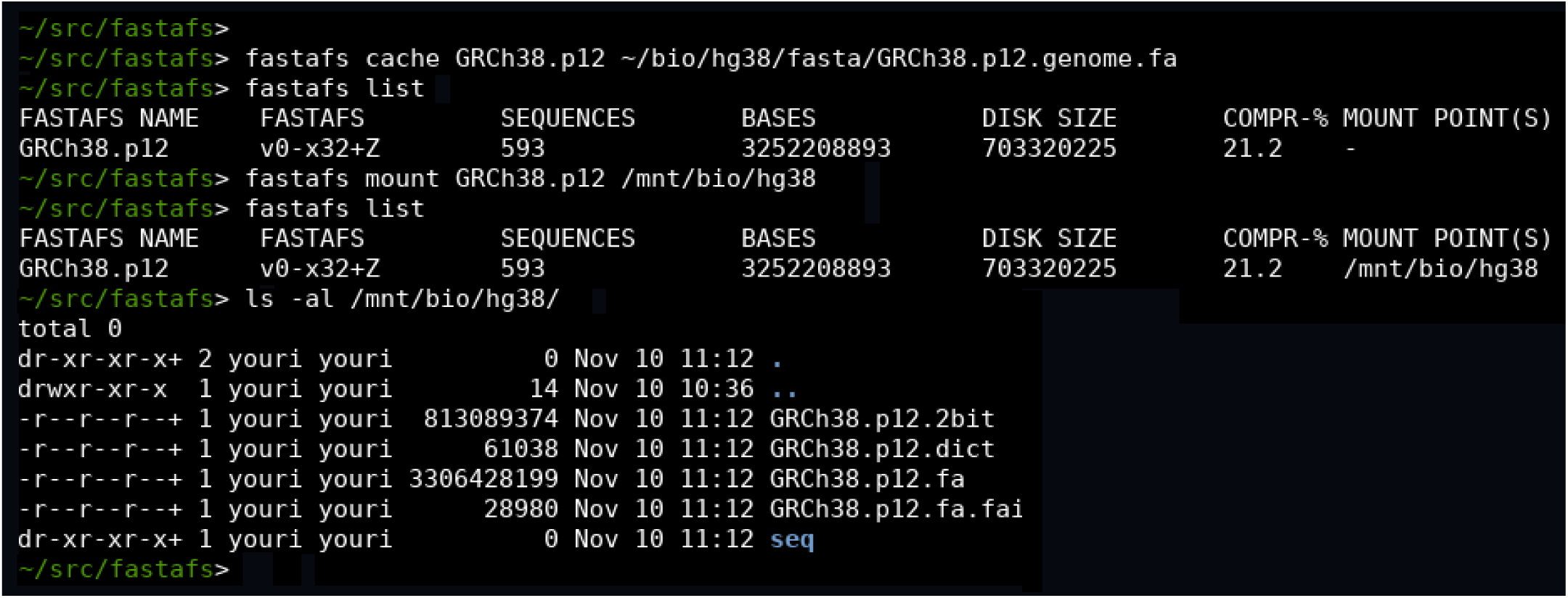
fastafs cache, mount & list. Screenshot of several `fastafs` commands: it starts by creating an archive using `fastafs cache`, followed by requesting the archives present on the system with `fastafs list`. It then mounts the archive to a mount point using `fastafs mount`. When the archives present at the system are listed with `fastafs list` again, the active mount point is shown. When we perform a system directory listing (`ls`), the virtual files and sizes are shown.

**Figure S1B:**
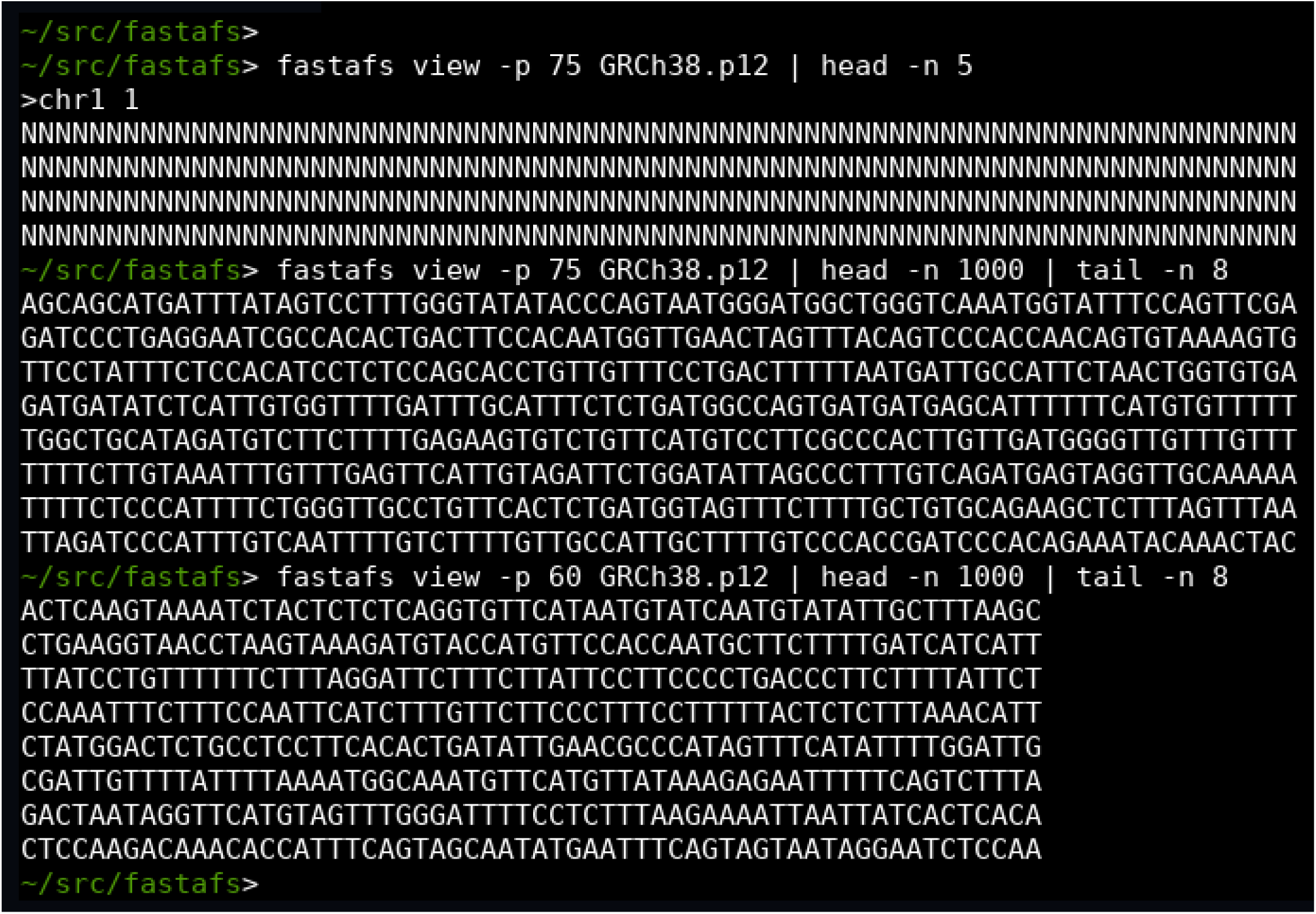
fastafs view. The `fastafs view` command writes directly to *stdout*. The padding size can be controlled with the `-p` argument.

**Figure S1C:**
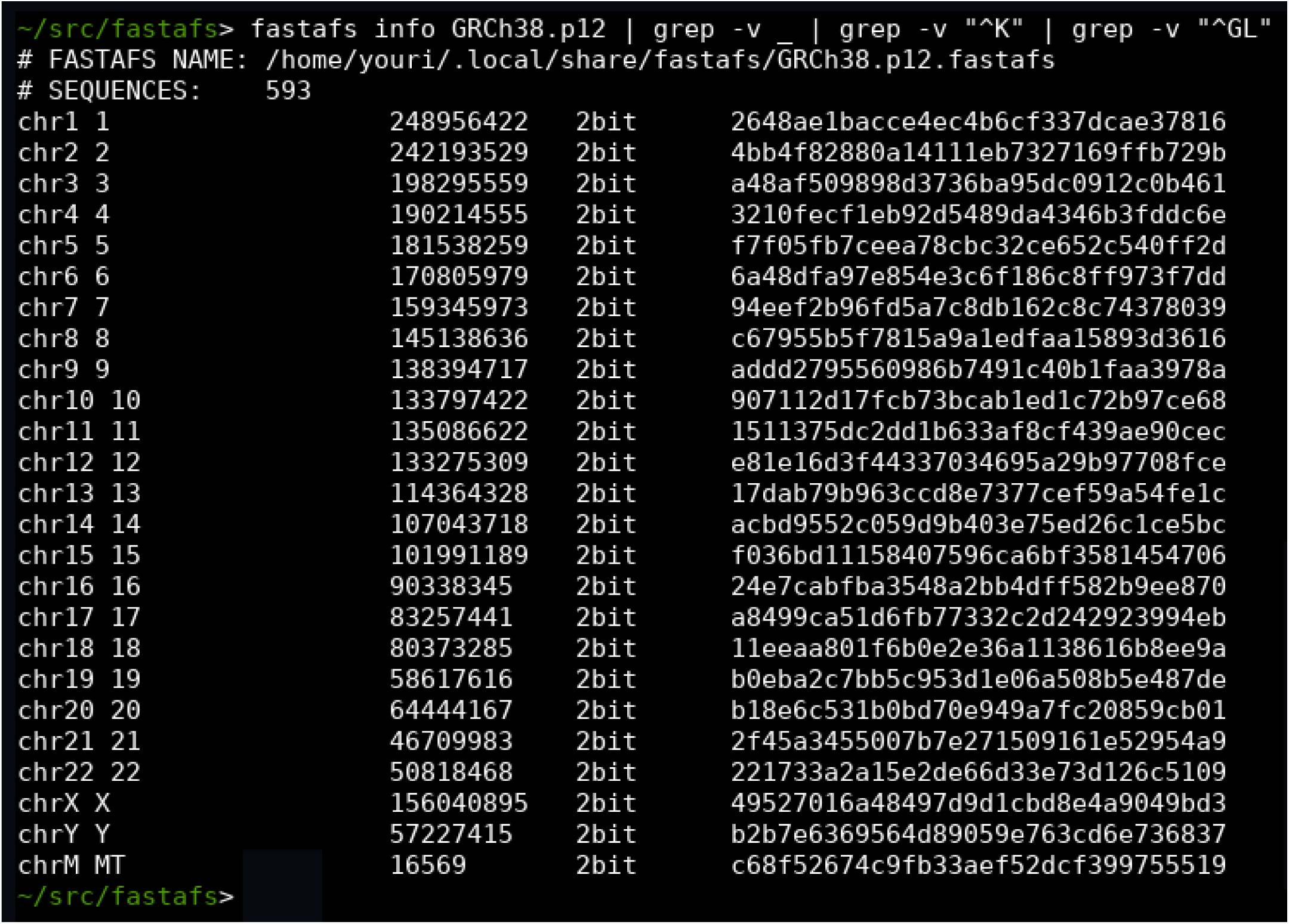
fastafs info. The command `fastafs info` shows general and per-sequence information for a given archive. The ENA compatible md5 checksums are provided in the last column.

**Figure S1D:**
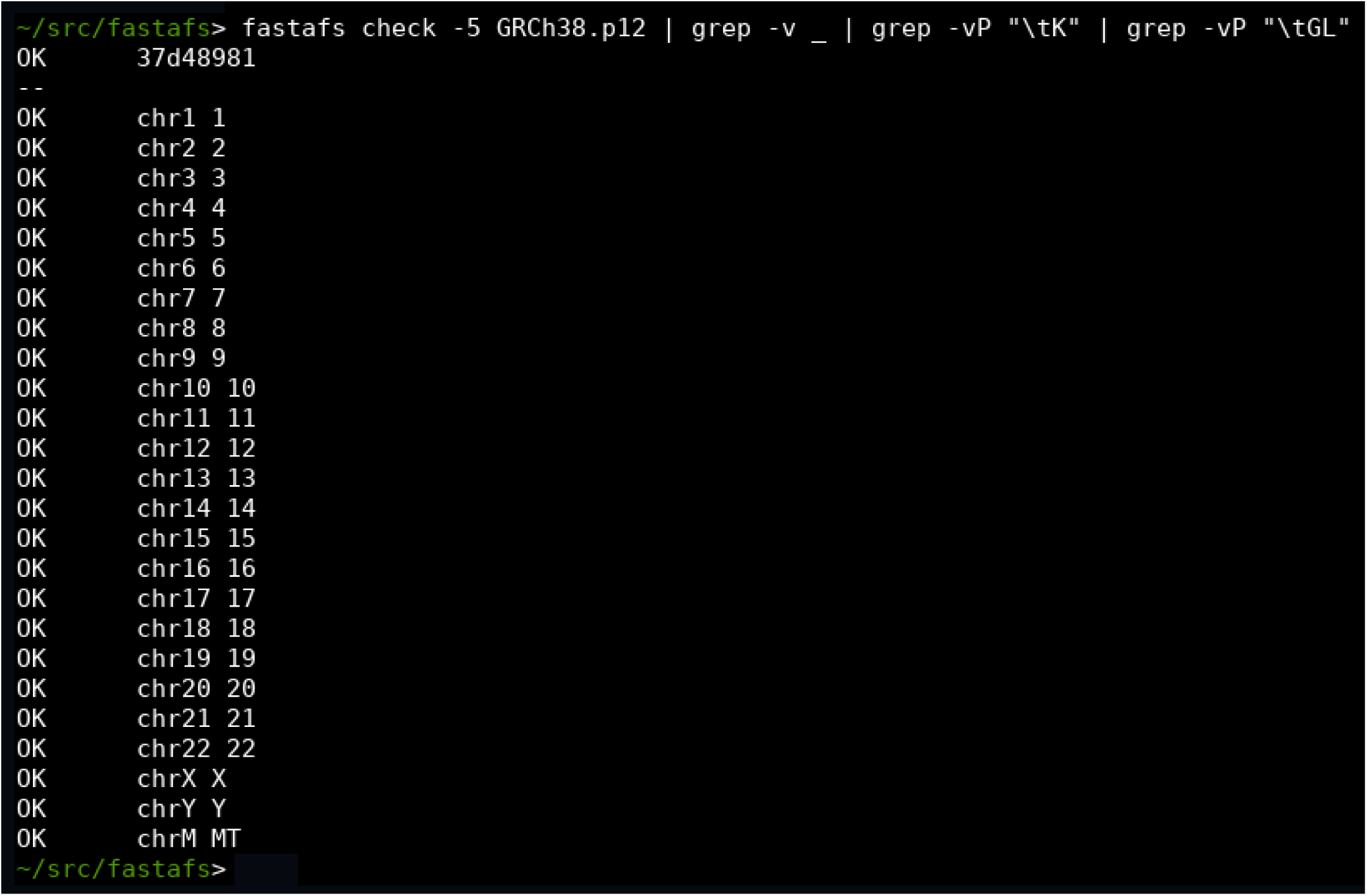
fastafs check. The `fastafs check` command checks the file integrity using a crc32 checksum. Using the optional `-5` argument, the per-sequence md5 checksum can be verified as well.

**Figure S1E:**
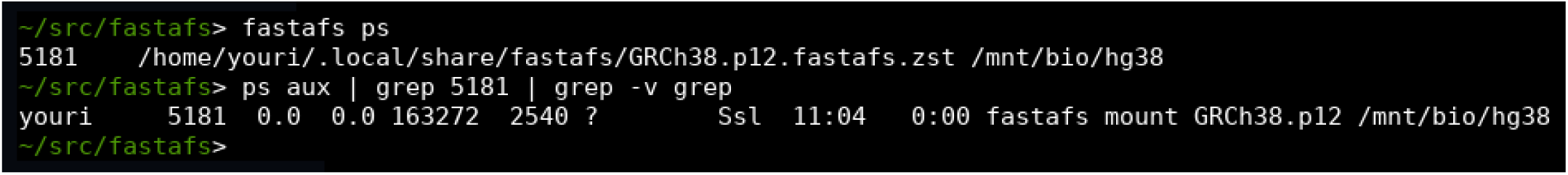
fastafs ps. The `fastafs ps` command can be used to retrieve all running instances of FASTAFS with corresponding process id’s and mount points.

